# SnapHiC: a computational pipeline to map chromatin contacts from single cell Hi-C data

**DOI:** 10.1101/2020.12.13.422543

**Authors:** Miao Yu, Armen Abnousi, Yanxiao Zhang, Guoqiang Li, Lindsay Lee, Ziyin Chen, Rongxin Fang, Jia Wen, Quan Sun, Yun Li, Bing Ren, Ming Hu

**Author notes:** Contributed equally (Miao Yu, Armen Abnousi).

## Abstract

Single cell Hi-C (scHi-C) analysis has been increasingly used to map the chromatin architecture in diverse tissue contexts, but computational tools to define chromatin contacts at high resolution from scHi-C data are still lacking. Here, we describe SnapHiC, a method that can identify chromatin loops at high resolution and accuracy from scHi-C data. We benchmark SnapHiC against HiCCUPS, a common tool for mapping chromatin contacts in bulk Hi-C data, using scHi-C data from 742 mouse embryonic stem cells. We further demonstrate its utility by analyzing single-nucleus methyl-3C-seq data from 2,869 human prefrontal cortical cells. We uncover cell-type-specific chromatin loops and predict putative target genes for non-coding sequence variants associated with neuropsychiatric disorders. Our results suggest that SnapHiC could facilitate the analysis of cell-type-specific chromatin architecture and gene regulatory programs in complex tissues.

## Main text

Transcriptional regulatory elements communicate with each other dynamically in the 3D nuclear space to direct cell-type-specific gene expression during development^1–3^. Understanding the transcriptional regulatory programs requires a high resolution view of the 3D chromatin architecture in the cell. Technologies have been developed to map chromatin architecture in single cells to explore the heterogeneity of chromatin organization in complex tissues^4–13^. However, it is still challenging to identify chromatin loops at the necessary resolution to delineate spatial proximity between transcriptional regulatory elements due to the extreme sparsity of the single cell chromatin contact matrix. The current strategy to identify chromatin loops from aggregated single cell Hi-C data from the same cell type with existing loop calling methods^14–18^ requires a large number of cells (>500-1,000), which is both cost prohibitive and impractical for the rare cell types in a complex tissue. Simulation studies^19^ showed that the sensitivity of existing loop calling methods decays exponentially with the decrease in the number of contacts. Here, we report single nucleus analysis pipeline for Hi-C (SnapHiC), a new computational framework that fully exploits the power of single cell Hi-C (scHi-C) data to identify chromatin loops at high resolution and accuracy.

SnapHiC identifies chromatin loops at 10-kilobase (Kb) resolution from scHi-C data by maximizing the usage of information from each single cell (**Fig. 1a** and **Methods**). Specifically, SnapHiC first imputes chromatin contact probability between all intra-chromosomal bin pairs with the random walk with restart (RWR) algorithm^20^ in each individual cell. Next, it converts the imputed contact probability into the normalized contact probability stratified based on linear genomic distances. SnapHiC then applies the paired *t*-test using all cells to identify loop candidates (see details in **Methods**). To remove false positives, SnapHiC considers a bin pair as a loop candidate only when it has significantly higher normalized contact probability than expected by chance based on both the global background and the local background. Finally, SnapHiC groups the loop candidates into discrete clusters using the Rodriguez and Laio’s algorithm^21^, and identifies the summit(s) within each cluster.

**Figure 1.**
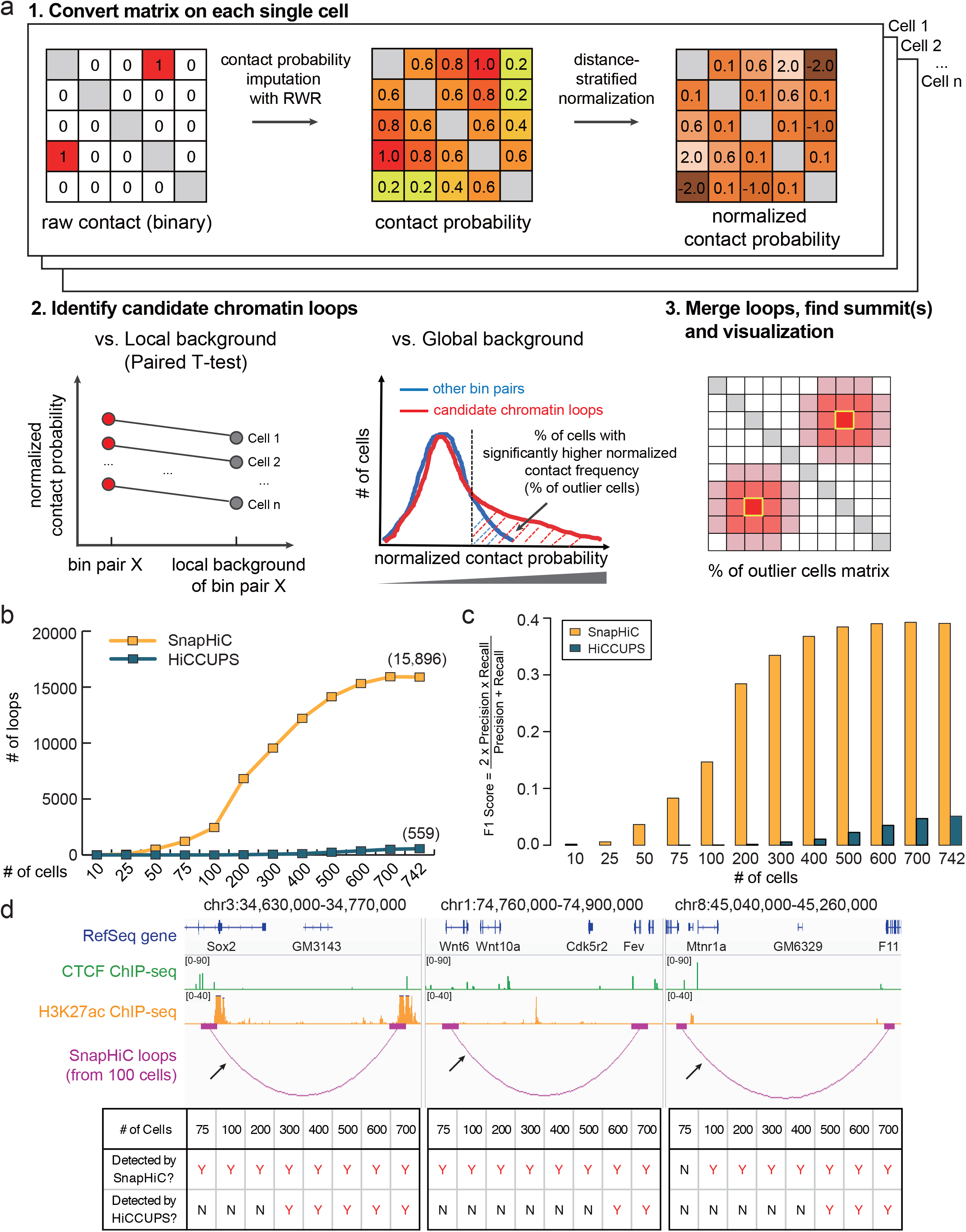
SnapHiC reveals chromatin loops at high resolution and accuracy. **(a)** Overview of SnapHiC workflow. The first step of SnapHiC is to convert the binary contact matrix to normalized contact frequency for each individual cell. Next, SnapHiC applies the paired *t*-test to identify candidate chromatin loops by comparing the normalized contact frequency of any given bin pair with its local and global background. Finally, SnapHiC merges nearby candidate loops into clusters and identifies the summit(s). Due to the sparsity of the raw count matrix of scHi-C data, the SnapHiC-identified loops can be visualized by the percentage of the outlier cells matrix. **(b)** The number of chromatin loops at 10Kb resolution identified by SnapHiC and HiCCUPS from different numbers of mES cells. **(c)** F1 score (the harmonic mean of the precision and recall) of SnapHiC- and HiCCUPS-identified loops from different numbers of mES cells. **(d)** (Top) Chromatin loops around *Sox2* (left)*, Wnt6* (middle), and *Mtnr1a* (right) gene identified from 100 mES cells using SnapHiC at 10Kb resolution. The black arrow points to the interaction verified in the previous publications^27,28^ with CRISPR/Cas9 deletion or 3C-qPCR. (Bottom) Comparison of the performance of SnapHiC and HiCCUPS (applied on aggregated scHi-C data) from the different number of mES cells at these three regions. If the previously verified interaction (black arrow) is recaptured, it is labeled as “Y”; otherwise, it is labeled as “N”.

To benchmark the performance of SnapHiC against a commonly used method, HiCCUPS^14^ designed for bulk Hi-C data analysis, we applied it to the published scHi-C data^5^ generated from mouse embryonic stem (mES) cells. We sub-sampled 10, 25, 50, 75, 100, 200, 300, 400, 500, 600, 700 and 742 cells from this dataset, and determined the intra-chromosomal loops at 10Kb resolution from 100Kb to 1Mb range. For each sub-sampling, we also pooled the scHi-C data and identified chromatin loops at 10Kb resolution within the same distance range using HiCCUPS. For each sub-sampling dataset, SnapHiC found more chromatin loops than HiCCUPS, suggesting that SnapHiC has a much higher sensitivity than HiCCUPS (**Fig. 1b** and **Supplementary Table 1-3**). Even from 75 cells, SnapHiC identified 1,219 loops, whereas HiCCUPS found only 2 loops. Additionally, HiCCUPS-identified loops tended to be a subset of SnapHiC-identified loops. For example, SnapHiC and HiCCUPS identified 15,896 and 559 loops from 742 cells, respectively, and 511 (91.4%) of HiCCUPS-identified loops are re-captured by SnapHiC (**Supplementary Table 1**). Moreover, SnapHiC achieves higher reproducibility than HiCCUPS for loop calling between replicates (for each replicate with 371 cells, 50.8% vs. 38.7%, paired *t*-test p-value = 7.86e-8, see details in **Methods**).

We used the F1 score, the harmonic mean of the precision and recall, to evaluate the overall performance of each method (see details in **Methods**). To calculate the F1 score, we combined long-range chromatin interactions identified by HiCCUPS from bulk *in situ* Hi-C data^22^, with interactions identified by MAPS from H3K4me3 PLAC-seq data^23^, cohesin^24^ and H3K27ac HiChIP data^25^, all from mES cells as a reference loop list (**Supplementary Table 4**). At each sub-sampling of scHi-C data, SnapHiC consistently attained a greater F1 score than HiCCUPS (**Fig. 1c, Supplementary Fig. 1**). The reliability of SnapHiC-identified loops can be further supported by two additional lines of evidence: 1) Significantly focal enrichment can be observed from aggregate peak analysis (APA) plots of SnapHiC-identified loops from the different number of cells (except for 10 cells) on aggregated scHi-C contact matrix from 742 cells (**Supplementary Fig. 2**); 2) For the SnapHiC-identified loops that have CTCF binding on both ends, there is a clear preference in convergent orientation – ranging from 63.6% to 78.7% when at least 50 cells are used for loop calling (**Supplementary Table 5**), as predicted by the loop extrusion model^14,26^.

The advantage of SnapHiC is more obvious when the number of cells profiled is limited. As illustrated in **Fig. 1d** (see also **Supplementary Fig. 3**), SnapHiC detected previously verified long-range interactions at *Sox2*, *Wnt6*, and *Mtnr1a* loci^27,28^ with as few as 75 or 100 cells, whereas HiCCUPS required at least 300-600 cells to detect the same loops. Taken together, the above results suggest that SnapHiC allows for the identification of chromatin loops from a small number of cells with high sensitivity and accuracy, underlining its potential utility in scHi-C data generated from complex tissues.

To demonstrate the utility of SnapHiC for analysis of scHi-C data from complex tissues, we applied SnapHiC to the published single-nucleus methyl-3C-seq (sn-m3C-seq) data^13^ from human prefrontal cortex, which simultaneously profiled DNA methylome and chromatin organization from the same cells. In this study, 14 major cell types were identified using CG and non-CG methylation. We applied SnapHiC to each of the 14 cell clusters and identified 817 ~ 27,379 loops at 10Kb resolution (**Fig. 2a** and **Supplementary Table 6**). Consistent with our observation on mES cells, SnapHiC identified more loops than HiCCUPS for all cell clusters, and more than 78% of HiCCUPS-identified loops are captured by SnapHiC (**Supplementary Table 7-8**). Except for oligodendrocytes, which have >1,000 cells, SnapHiC found ~4-70 folds more loops than HiCCUPS in other 13 cell types. We also calculated the F1 scores of SnapHiC- and HiCCUPS-identified chromatin loops in oligodendrocytes, microglia, and eight neuronal subtypes, and benchmarked against promoter-centered chromatin contacts previously identified from H3K4me3 PLAC-seq analysis of purified oligodendrocytes, microglia, astrocytes and neurons (**Supplementary Table 9**)^29^. Again, SnapHiC achieved much greater F1 scores than HiCCUPS in each cell cluster (**Fig. 2b** and **Supplementary Fig. 4**).

**Figure 2.**
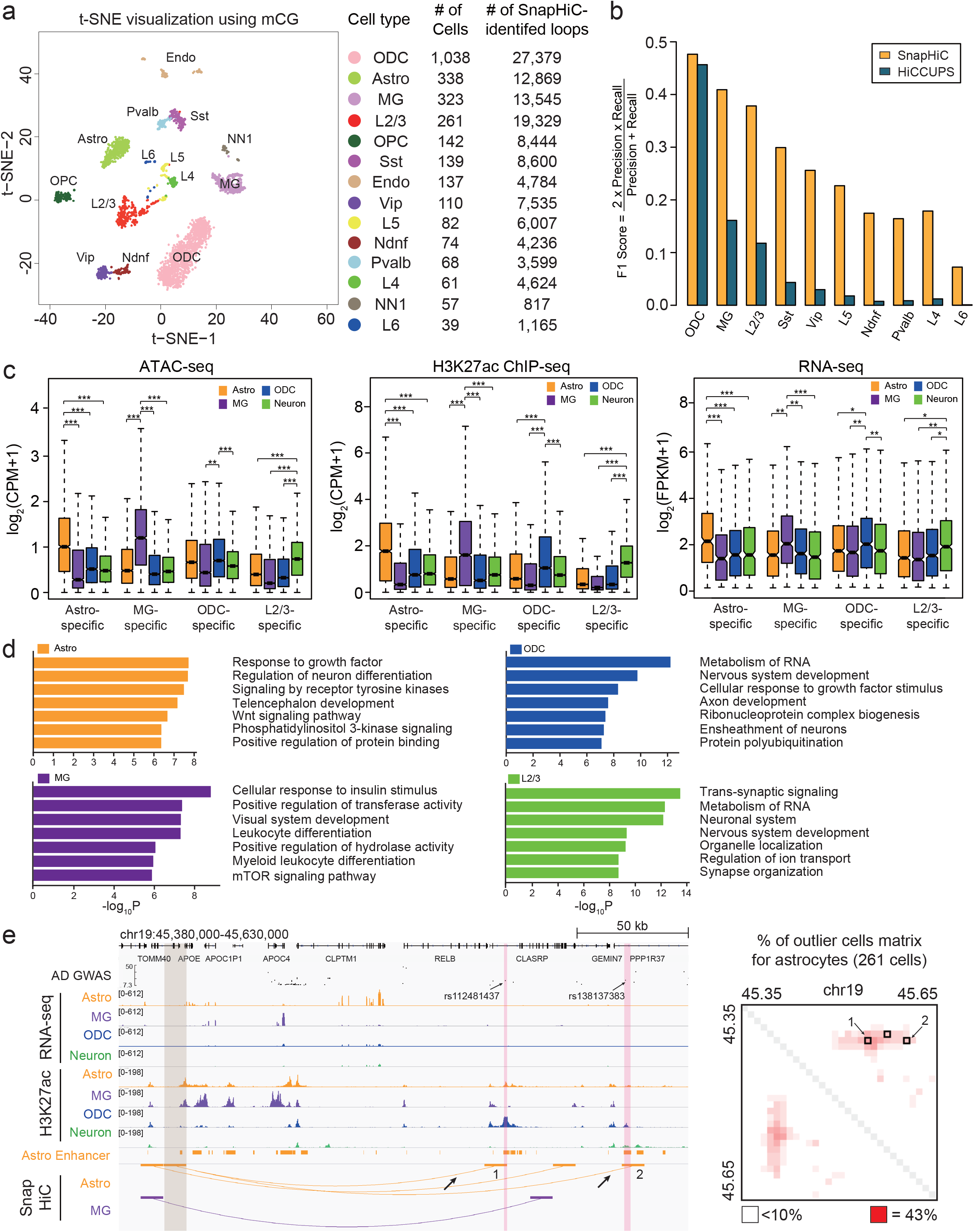
Application of SnapHiC to sn-m3C-seq data from human prefrontal cortex uncovered chromatin loops in diverse brain cell types. **(a)** (Left) t-SNE visualization of 14 major cell types identified in human prefrontal cortex in Lee et al. study^13^ using CG methylation of non-overlapping 100Kb genomic bins. ODC: oligodendrocyte. Astro: astrocyte. MG: microglia. OPC: oligodendrocyte progenitor cell. Endo: endothelial cell. L2/3, L4, L5 and L6: excitatory neuron subtypes located in different cortical layers. Pvalb and Sst: medial ganglionic eminence-derived inhibitory subtypes. Ndnf and Vip: CGE-derived inhibitory subtypes. NN1: non-neuronal cell type 1. (Right) The number of cells and SnapHiC-identified loops in each of the 14 cell types. **(b)** F1 score (the harmonic mean of the precision and recall) of SnapHiC- and HiCCUPS-identified loops for oligodendrocytes (ODC), microglia (MG) and eight neuronal subtypes. **(c)** Boxplot of ATAC-seq log2(CPM+1) value (left), H3K27ac ChIP-seq log2(CPM+1) value (middle) and RNA-seq log_2_(FPKM+1) value (right) in astrocyte, microglia, oligodendrocytes and neurons at the anchors of Astro-specific, MG-specific, ODC-specific, L2/3-specific SnapHiC loops summarized in **Supplementary Table 11**. \*\*\**p* < 2.2e-16; \*\**p* < 1e-10; \**p* < 1e-7 by the paired Wilcoxon signed-rank test. **(d)** Top seven enriched gene ontology (GO) terms of genes associated with cell-type-specific SnapHiC loops. **(e)** (Left) SnapHiC-identified loops from astrocyte and microglia around gene *APOE*. There is no loop identified in this genomic region from oligodendrocytes or L2/3 excitatory neurons, so no corresponding tracks are shown. Two astrocyte-specific loops linking the *APOE* promoter (highlighted in grey) and the active enhancers in astrocyte (highlighted in pink) containing two AD-associated GWAS SNPs are marked by black arrows. Only *APOE* TSS-distal AD-associated GWAS SNPs are shown in the figures (residing in the region chr19: 45,440,000-45,630,000). (Right) Matrix of the percentage of cells with significantly higher normalized contact frequency (percentage of outlier cells with normalized contact frequency>1.96) for 261 astrocytes. The SnapHiC-identified loops from astrocyte are marked by black squares.

The accuracy and sensitivity of SnapHiC are further supported by several lines of evidence. First, APA analysis confirms that SnapHiC-identified loops show significant enrichment of contacts compared to their local background on the aggregated contact matrix from cells in the corresponding cluster (**Supplementary Fig. 5**). Next, SnapHiC-identified loops correlate with cell-type-specific chromatin accessibility, histone acetylation, and gene expression. For this analysis, we focused on four distinct cell types, astrocytes, L2/3 excitatory neurons, oligodendrocytes and microglia, in which ATAC-seq, H3K27ac ChIP-seq and RNA-seq data are available^29,30^. To minimize the effect of cell number variation between different cell types, we randomly selected the same number of cells (N=261) from astrocytes, oligodendrocytes and microglia to match the number of cells available from L2/3 excitatory neurons, and applied SnapHiC to identify loops from these sub-sampled data (**Supplementary Table 10**). We found that most chromatin loops are cell-type-specific (**Supplementary Table 11,** see details in **Methods**). Further analysis showed that the anchors of cell-type-specific loops show significantly higher ATAC-seq and H3K27ac ChIP-seq signals in the matched cell type compared to those in the other three cell types (**Fig. 2c**). In addition, we found 407, 616, 860 and 1,002 genes whose promoters link to astrocyte-, microglia-, oligodendrocyte- and L2/3 excitatory neurons-specific loops, respectively (**Supplementary Table 12**). These genes show significantly higher expression levels in the matched cell type than those in the other three cell types (**Fig. 2c**) and are associated with gene ontology terms^31^ related to cell-type-specific biological processes (**Fig. 2d**). Taken together, our results suggest that SnapHiC can detect chromatin contacts reliably from single cell Hi-C data in complex tissues.

How sequence variations determine the phenotypic traits and propensity to human diseases is one of the fundamental questions in biology^32^. It is generally believed that many disease-associated non-coding variants contribute to disease etiology by perturbing the transcriptional regulatory sequences and affecting target gene expression^33–35^. The current catalogs of genes and candidate regulatory sequences in the human genome^33–37^ still lack the information about the target genes of annotated candidate *cis-*regulatory elements, making it a challenge to interpret the biological roles of non-coding risk variants. We used SnapHiC-identified loops in the four brain cell types (astrocytes, microglia, oligodendrocytes and L2/3 excitatory neurons) to assign candidate target genes to non-coding GWAS SNPs. We first collected 30,262 genome-wide significant (*p*-value<5e-8) non-coding GWAS SNP-trait associations from seven neuropsychiatric disorders and traits, including Alzheimer’s diseases^38^ (AD), attention deficit hyperactivity disorder^39^ (ADHD), autism spectrum disorder^40^ (ASD), bipolar disorder^41^ (BIP), intelligence quotient^42^ (IQ), major depressive disorder^43^ (MDD) and schizophrenia^44^ (SCZ), resulting in a total of 28,099 unique GWAS SNPs (**Supplementary Table 13**). We then focused on 3,639 SNP-disease associations (3,471 unique GWAS SNPs), where the corresponding SNPs reside within active enhancers of astrocytes, neurons, microglia or oligodendrocytes defined in the previous study^29^ (**Supplementary Table 13**). Using SnapHiC loops from the matching cell types (L2/3 excitatory neurons to represent neurons, all four cell types with 261 cells), we found 788 SNP-disease-loop-gene linkages, connecting 445 SNP-disease associations (416 unique GWAS SNPs) to 189 genes via 175 loops (**Supplementary Table 14**). Notably, such a list of GWAS SNP-interacting genes includes several known disease risk genes, including *APOE* (AD), *GRIN2A* (IQ), *INPP5D* (AD), *RAB27B* (MDD), *SORL1* (AD), *THRB* (IQ), and *ZNF184* (SCZ and MDD). **Fig. 2e** shows an illustrative example of gene *APOE*, which is specifically expressed in astrocyte. We found two astrocyte-specific chromatin loops, connecting the TSS of *APOE* to two active enhancers in astrocyte, ~150Kb and ~200Kb downstream, respectively. These two enhancers also contain two AD-associated GWAS SNPs, rs112481437 and rs138137383. Our data suggest that *APOE* is the putative target gene of these two GWAS SNPs only in astrocytes.

In summary, we describe SnapHiC, a novel method customized for sparse single cell Hi-C datasets to identify chromatin loops at high resolution and accuracy. Re-analysis of published single cell Hi-C data from mES cells demonstrate that SnapHiC greatly boosts the statistical power in loop detection. Application of SnapHiC to sn-m3C-seq data from human prefrontal cortical cells reveals cell-type-specific loops, which can be used to predict putative target genes of non-coding GWAS SNPs. SnapHiC has the potential to facilitate the study of cell-type-specific chromatin spatial organization in complex tissues.

## Code availability

SnapHiC software package with a detailed user tutorial and sample input and output files can be found at: https://github.com/HuMingLab/SnapHiC.

## Acknowledgements

We thank 4D Nucleome consortium investigators for comments and suggestions on the early version of this work. This study was funded by U54DK107977, UM1HG011585 (to B.R. and M.H.), and U01DA052713, R01GM105785 and P50HD103573 (to Y.L.).

## Author Contributions

This study was conceived and designed by M.H. and B.R.; Data analysis was performed by M.H., M.Y., A.A., Y.Z., G.L., L.L., Z.C., R.F., J.W., Q.S. and Y.L.; SnapHiC software package was developed by A.A. and M.H.; Manuscript was written by M.H., M.Y. and B.R. with input from all authors.

---

**Supplementary Figure 1.**
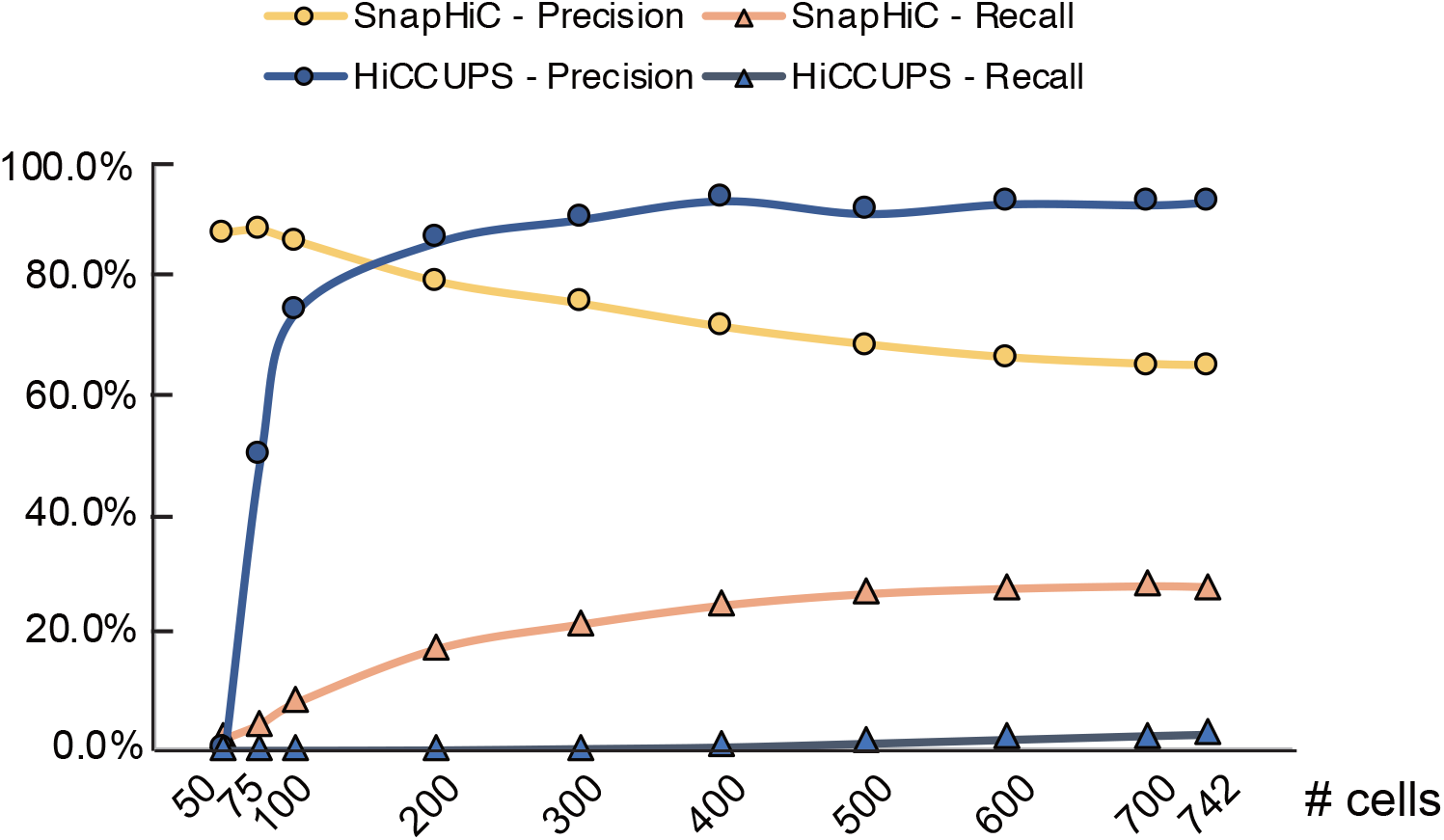
Comparison of the precision and recall values of SnapHiC- and HiCCUPS-identified loops from mES cells. The precision and recall values are calculated for the loops identified by SnapHiC and HiCCUPS from different numbers of mES cells. These values are also used to calculate the F1 score in **Fig. 1c**.

**Supplementary Figure 2.**
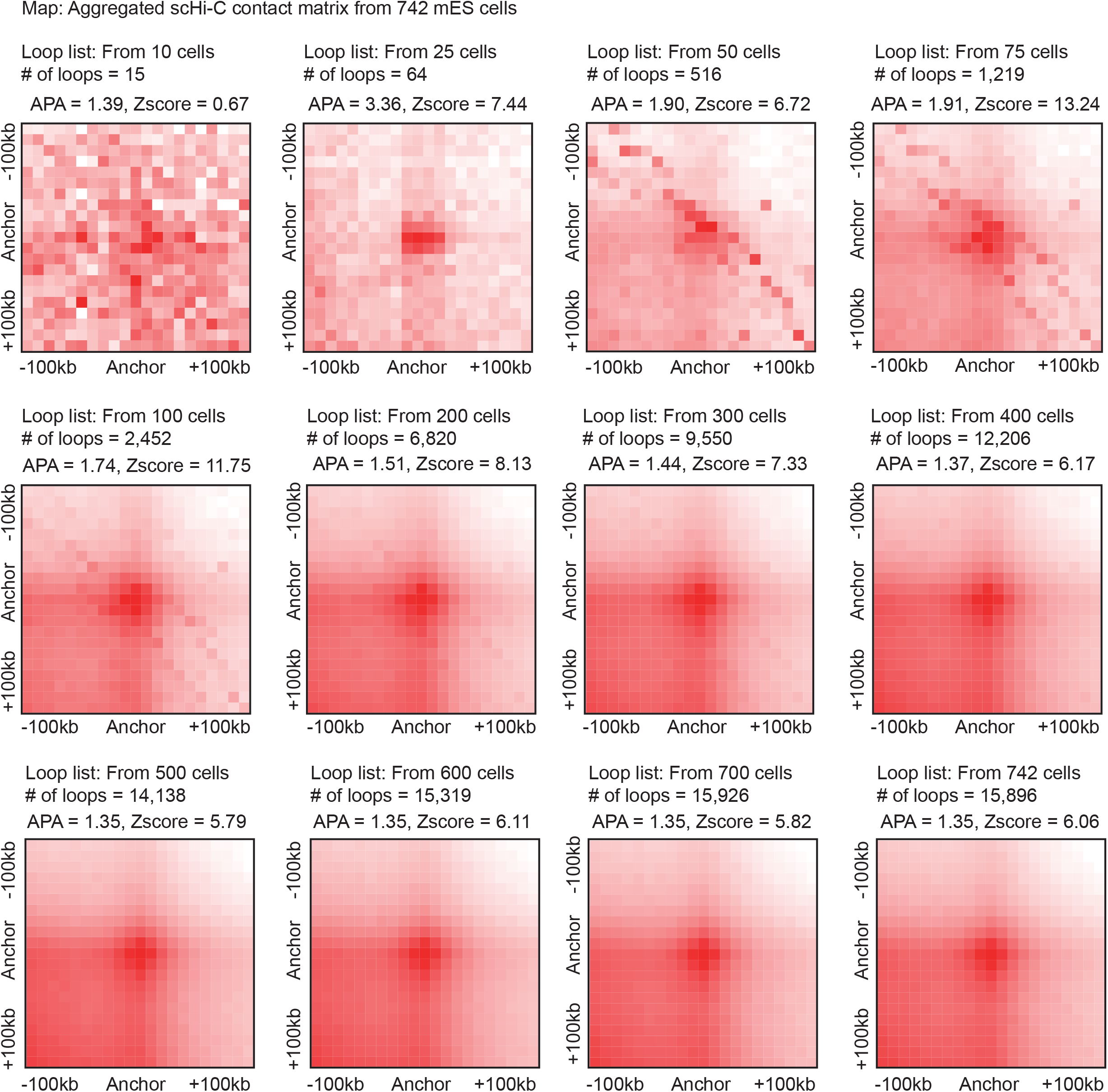
SnapHiC-identified loops from different sub-sampling of mES cells show significant enrichment over their local background. Aggregate peak analysis (APA) of SnapHiC-identified loops from different sub-sampling of mES cells examined on aggregated scHi-C contact matrix of 742 cells.

**Supplementary Figure 3.**
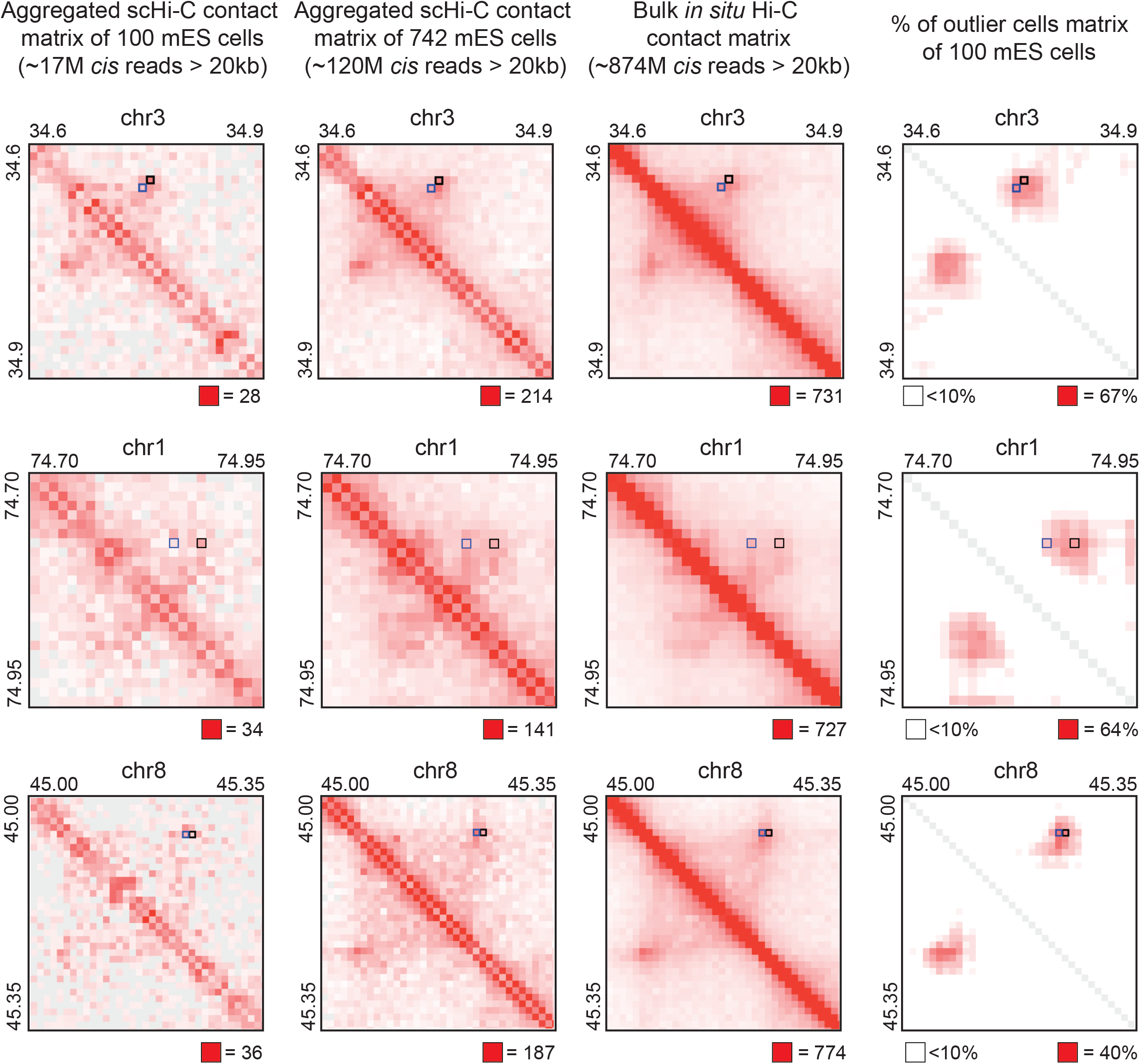
Visualization of selected SnapHiC-identified loops. From left to right: aggregated scHi-C contact matrix of 100 mES cells, aggregated scHi-C contact matrix of 742 mES cells, bulk *in situ* Hi-C contact matrix from mES cells (replicate 1 from Bonev et al. study^22^) and % of outlier cells matrix of 100 mES cells at 10Kb resolution; from top to bottom: *Sox2* locus, *Wnt6* locus, and *Mtnr1a* locus. Black squares represent the SnapHiC-identified loops from 100 mES cells, which are shown in **Fig. 1d** as purple arcs. For comparison, the HiCCUPS-identified loops from the deepest available bulk *in situ* Hi-C data of mES cells (combining all four replicates from Bonev et al. study^22^) are marked as blue squares.

**Supplementary Figure 4.**
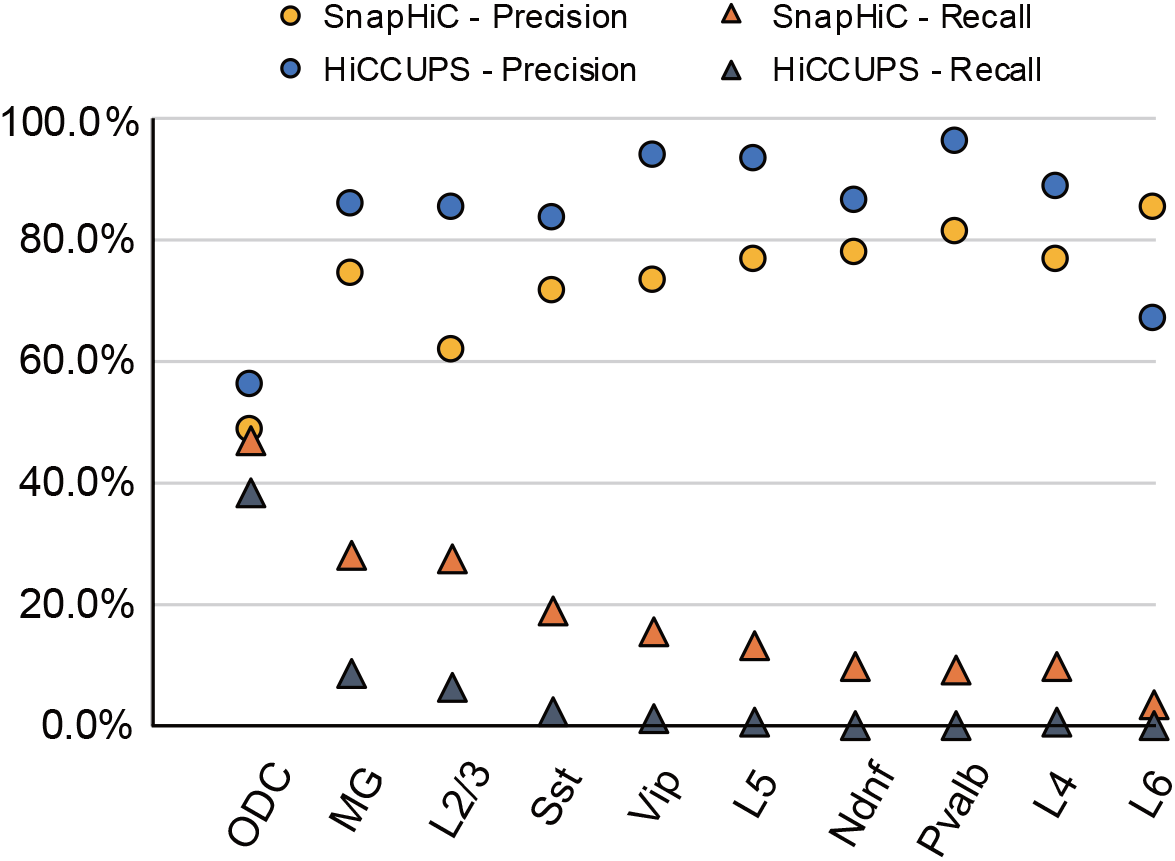
Comparison of the precision and recall values of SnapHiC- and HiCCUPS-identified loops for ten cell clusters from human prefrontal cortex. The precision and recall values are calculated for the loops identified by SnapHiC and HiCCUPS for oligodendrocytes (ODC), microglia (MG), and eight neuronal subtypes. These values are also used to calculate the F1 score in **Fig. 2b**.

**Supplementary Figure 5.**
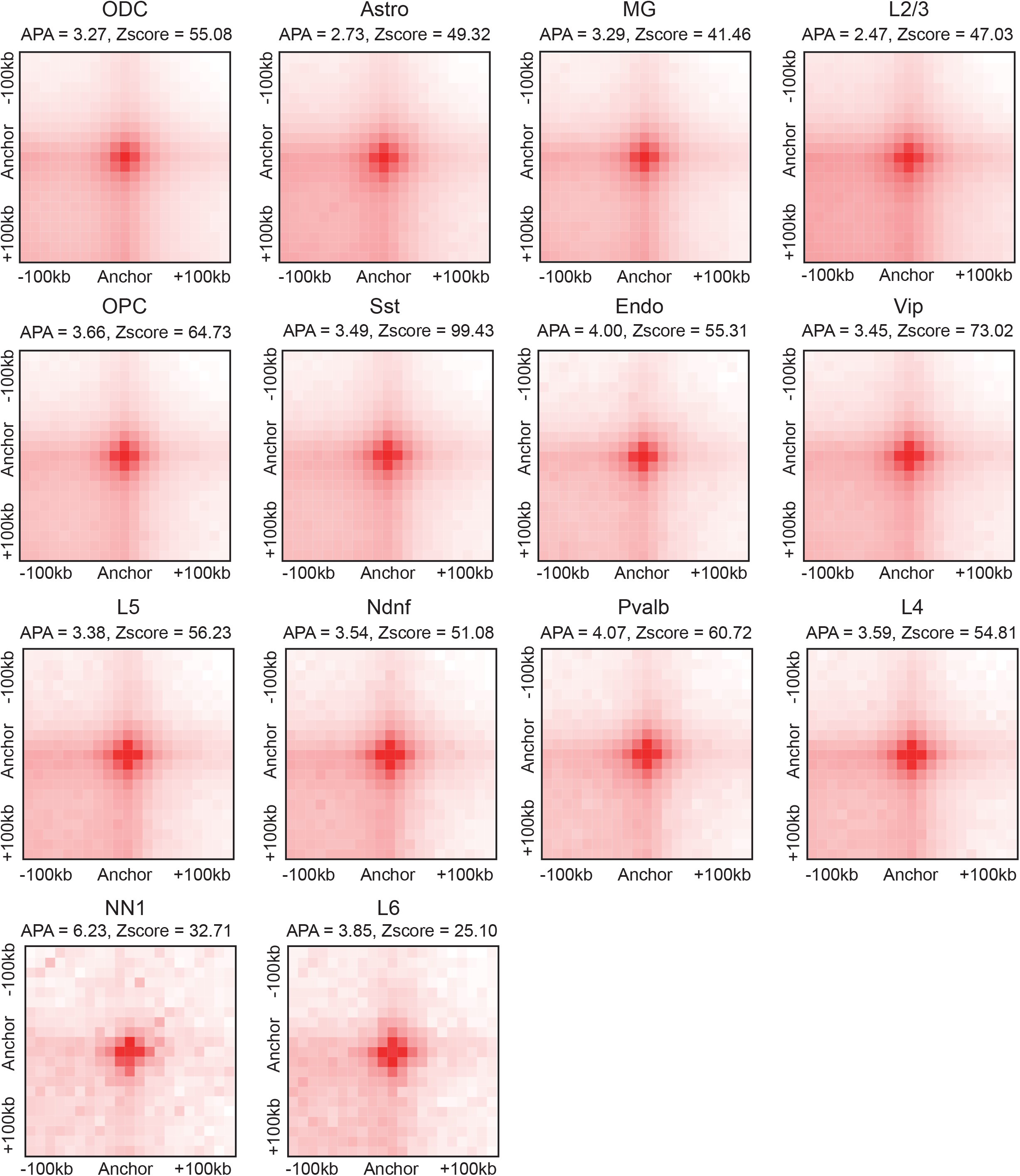
SnapHiC-identified loops from each of the 14 cell clusters identified from sn-m3C-seq data of the human prefrontal cortex show significant enrichment over their local background. Aggregate peak analysis (APA) of SnapHiC-identified loops for each of the 14 cell clusters demonstrated in **Fig. 2a** examined on the aggregated contact matrix from the matching cell clusters.

**Supplementary Figure 6.**
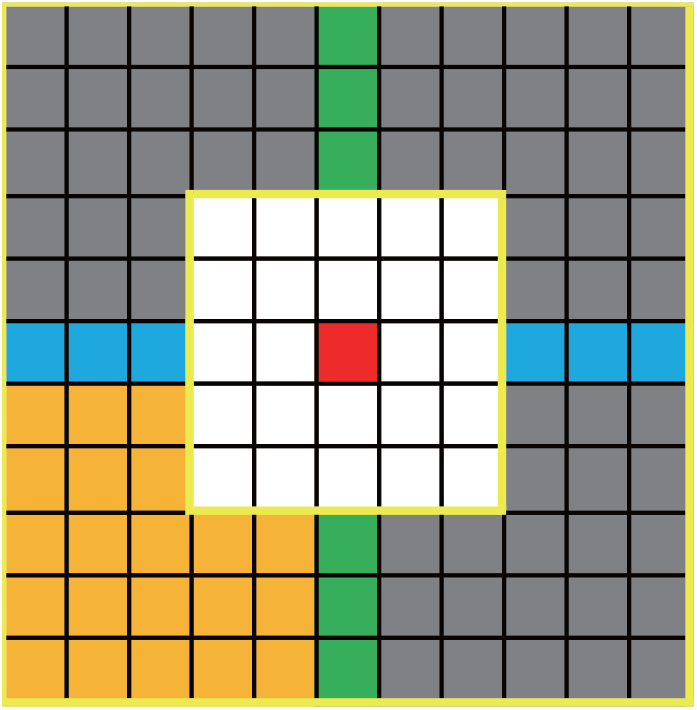
Illustration of different types of the local background used for SnapHiC loop calling. For each 10Kb bin pair of interest (red), its horizontal background, vertical background, lower left background and donut background are the blue, green, yellow and grey areas, respectively. The circle background, which is also the local neighborhood, is the union of the blue, green, yellow and grey areas.

**Supplementary Figure 7.**
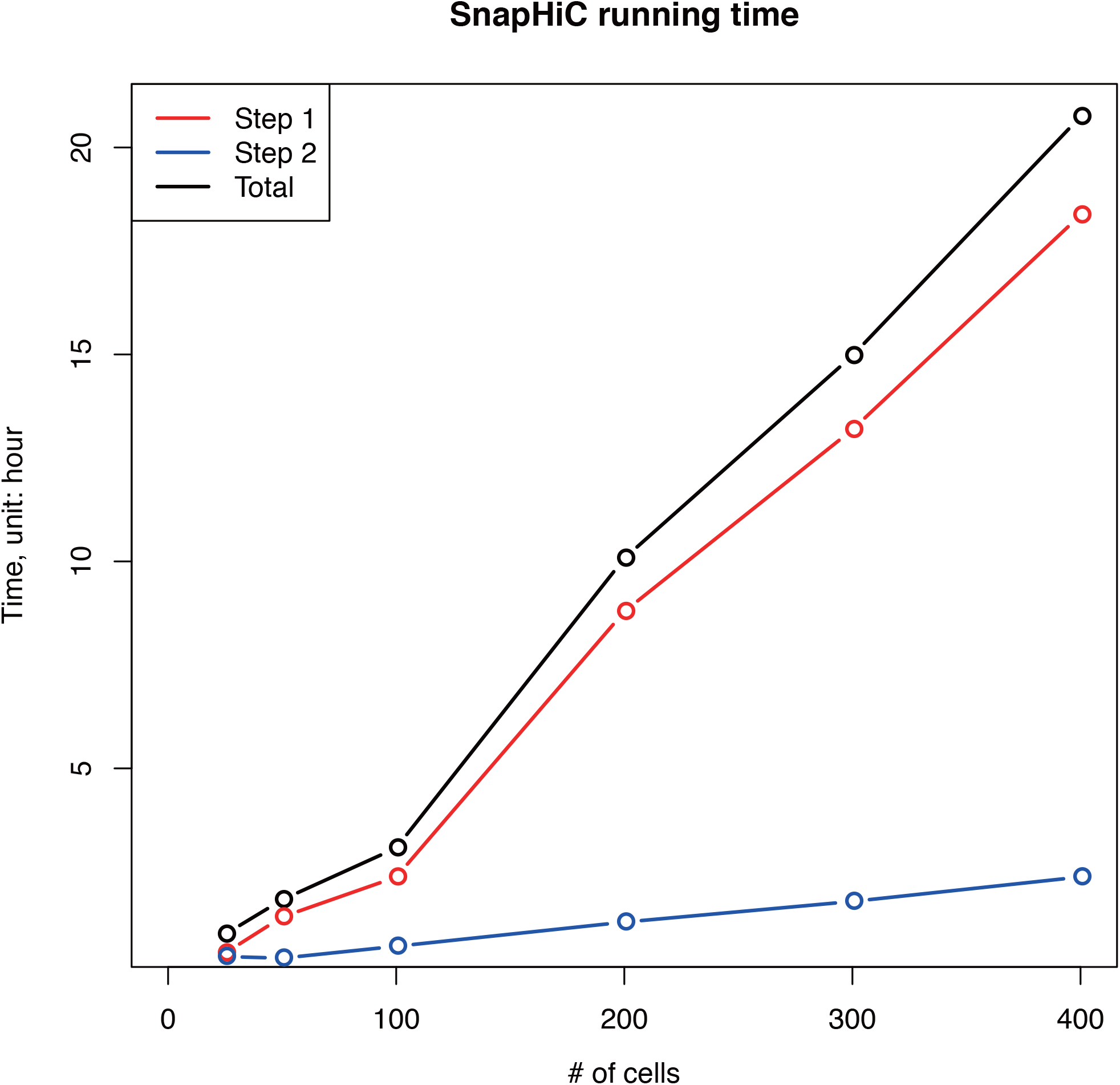
The relationship between the number of cells and the running time of SnapHiC analysis. We tested the running time of SnapHiC on scHi-C data from 25, 50, 100, 200, 300 and 400 mES cells (10Kb resolution, searching for loops between 100Kb to 1Mb genomic distance). SnapHiC consists of two steps: (1) applying the random walk with restart (RWR) algorithm to impute contact probability within every single cell, and (2) integrating imputed contact probability matrices from all single cells to identify chromatin loops. The running time of each step and the sum of both steps against the number of cells is plotted.

## Supplementary Table Legends

**Supplementary Table 1.** Summary of SnapHiC- and HiCCUPS-identified loops from mES scHi-C data. Related to **Fig. 1b**.

**Supplementary Table 2.** SnapHiC-identified loops from 10, 25, 50, 75, 100, 200, 300, 400, 500, 600, 700 and 742 mES cells (Column D in **Supplementary Table 1**).

**Supplementary Table 3.** HiCCUPS-identified loops from 10, 25, 50, 75, 100, 200, 300, 400, 500, 600, 700 and 742 mES cells (after filtering, Column F in **Supplementary Table 1**).

**Supplementary Table 4.** HiCCUPS-identified loops from bulk *in situ* Hi-C data, MAPS-identified significant interactions from H3K4me3 PLAC-seq, cohesin HiChIP, and H3K27ac HiChIP data, which are used as the reference loop list after pooling to calculate precision, recall values and the F1 score in **Fig. 1c** and **Supplementary Fig. 1**.

**Supplementary Table 5.** CTCF motif orientation analysis for SnapHiC-identified loops from different numbers of mES cells.

**Supplementary Table 6.** SnapHiC-identified loops from 14 different cell clusters demonstrated in **Fig. 2a** (Column D in **Supplementary Table 8**).

**Supplementary Table 7.** HiCCUPS-identified loops from 14 different cell clusters demonstrated in **Fig. 2a** (after filtering, Column F in **Supplementary Table 8**).

**Supplementary Table 8.** Summary of SnapHiC- and HiCCUPS-identified loops from 14 different cell clusters demonstrated in **Fig. 2a**.

**Supplementary Table 9.** MAPS-identified interaction lists of human microglia, oligodendrocytes, and neurons based on the Nott. et al. study^29^, which are used to calculate precision, recall values and the F1 score in **Fig. 2b** and **Supplementary Fig. 4**.

**Supplementary Table 10.** SnapHiC-identified loops from astrocytes, microglia and oligodendrocytes after sub-sampling (261 cells for each cell type).

**Supplementary Table 11.** Cell-type-specific SnapHiC loops identified from astrocytes, microglia, oligodendrocytes, and L2/3 excitatory neurons after sub-sampling (261 cells for each cell type).

**Supplementary Table 12.** Genes whose promoter overlaps with cell-type-specific SnapHiC loops identified from astrocytes, microglia, oligodendrocytes, and L2/3 excitatory neurons after sub-sampling (261 cells for each cell type). Related to **Fig. 2c**, **2d** and **Supplementary Table 11**.

**Supplementary Table 13.** Non-coding GWAS SNPs associated with seven neuropsychiatric disorders with *p*-value < 5×10^−8^ and the SNPs residing in the active enhancers of astrocytes, microglia, oligodendrocytes or neurons defined in the previous publication^29^.

**Supplementary Table 14.** Predicted 788 SNP-disease-loop-gene quadruplets using SnapHiC-identified loops in astrocytes, microglia, oligodendrocytes and L2/3 excitatory neurons (261 cells for each cell type).

## Methods

### Single-cell Hi-C (scHi-C) data processing

For scHi-C data from mES cells^5^, we downloaded the raw fastq files of all diploid serum cells (in total 1,175 cells). We first aligned scHi-C read pairs for each single cell to mm10 genome with BWA-MEM with the “-5” option to report the most 5’ end alignment as the primary alignment, and the “-P” option to perform Smith-Waterman algorithm to rescue chimeric reads. We only used primary alignments in the next steps. We then de-duplicated read pairs with the Picard tool to keep only one read pair at the exact same position. We further applied two filtering steps to remove read duplications: (1) we split each chromosome into consecutive non-overlapping 1Kb bins, and only kept one contact for each 1Kb bin pair, (2) we removed 1Kb bins which contact with more than 10 other 1Kb bins, since they are likely mapping artifacts. We found that the number of contacts per cell for these 1,175 cells has a bimodal distribution, therefore we selected the top 742 cells with >150,000 contacts per cell for downstream analysis.

### Single-nucleus methyl-3C-seq (sn-m3C-seq) data processing

For sn-m3C-seq data from human prefrontal cortex, we performed data processing using reference genome hg19 as described in the previous study^13^. After this processing, we also applied two additional filtering steps to remove read duplications as described in the “**Single-cell Hi-C (scHi-C) data processing**” section. Similar to scHi-C data from mES cells, we also observed a bimodal distribution in the number of contacts per cell for all 4,238 cells. Again, we selected the top 2,869 cells with >150,000 contacts per cell for downstream analysis. The method for clustering and cell type annotation for these 2,869 cells was the same as previously described^13^.

### SnapHiC algorithm

#### Step A. Contact probability imputation using the random walk with restart (RWR) algorithm

We first partitioned each autosomal chromosome into consecutive non-overlapping bins at a pre-specified resolution (10Kb in this study) and dichotomized contact for each 10Kb bin pair (binary contact matrix with 1 indicating non-zero contact and 0 otherwise). Next, we modeled each autosomal chromosome as an unweighted graph, where each 10Kb bin is one node, and each non-zero contact between any two 10Kb bins is one edge. We also added edges to all adjacent 10Kb bins. We then implemented the random walk with restart (RWR) algorithm^20^ with the restart probability 0.05 to impute the contact probability between all intra-chromosomal 10Kb bin pairs. We used the Python “NetworkX” package to construct the graph, and adopted the “linalg.solve” function in the Python “SciPy” package to solve the linear equation in the RWR algorithm. In addition, we distributed the analysis for different chromosomes in different cells between different processors using the Python “mip4py” package to speed up the computation.

We further evaluated whether the contact probability imputed by the RWR algorithm in each single cell contains systematic biases, including effective fragment size, GC content and mappability, which are known systematic biases in bulk Hi-C data^45^. Specifically, for each of the 742 mES scHi-C profiles, we used the RWR algorithm to impute the contact probability between all intra-chromosomal 10Kb bin pairs (*i*, *j*) within 1Mb genomic distance, denoted as *x*_*ij*_. Let *F*_*i*_, *GC*_*i*_ and *M*_*i*_ represent the effective fragment size, GC content and mappability of the 10Kb bin *i*, which are calculated according to our previous work^45^. We define *f*_*ij*_ = *F*_*i*_ * *F*_*j*_, *gc*_*ij*_ = *GC*_*i*_ * *GC*_*j*_, and *m*_*ij*_ = *M*_*i*_ * *M*_*j*_, as the measure of three types of bias for each 10Kb bin pair. We then calculated the Pearson Correlation Coefficient between the contact probability *x*_*ij*_ and *f*_*ij*_, *gc*_*ij*_ and *m*_*ij*_, respectively, for each of the 19 autosomal chromosomes in one cell. Next, we used the average Pearson Correlation Coefficient (aPCC) across all chromosomes as the measurement of bias in each cell. Among all 742 cells, the mean of aPCC is 0.0110, 0.0085 and −0.0016 for effective fragment size, GC content and mappability, respectively. The standard deviation of aPCC is 0.0068, 0.0113 and 0.0029 for effective fragment size, GC content and mappability, respectively. These results suggest that the systematic biases in imputed contact probabilities in scHi-C data are negligible, thus normalization against effective fragment size, GC content or mappability is not needed.

#### Step B. Contact probability normalization based on 1D genomic distance

Since the contact probability between any two genomic loci is strongly dependent on their 1D genomic distance, normalization of the imputed contact probability against 1D genomic distance is needed before loop calling. To achieve this, we first removed the bin pairs residing in the first 50Kb or the last 50Kb of each chromosome, which often have unusually high imputed contact probability due to the edge effect of the RWR algorithm. We then stratified all intra-chromosomal 10Kb bin pairs by their 1D genomic distance. Specifically, let *x*_*ij*_ represent the contact probability between bin *i* and bin *j*. Define the set *A*_*d*_ as all bin pairs (*i*, *j*) with the 1D genomic distance *d*. For simplicity, we only considered bin pairs (*i*, *j*) in the upper triangle of the contact matrix where *i* < *j*. We removed the top 1% bin pairs in *A*_*d*_ with the highest contact probability, and then computed the mean *μ*_*d*_ and the standard deviation *σ*_*d*_ of the contact probability using the remaining bin pairs in *A*_*d*_. We further calculated the normalized contact probability (i.e., Z-score), defined as *z*_*ij*_ = (*x*_*ij*_ − *μ*_*d*_)/*σ*_*d*_, for all bin pairs in *A*_*d*_. For single cells with very few contacts, the imputed contact probabilities *x*_*ij*_ at specific 1D genomic distance *d* are close to zero, leading to very small standard deviation *σ*_*d*_ and numerical errors in the Z-score transformation. To avoid this issue, when *σ*_*d*_ is less than 1e-6, we defined *z*_*ij*_ = 0 for all bin pairs in *A*_*d*_. After the calculation described above, bin pair (*i*, *j*) with higher normalized contact probability *z*_*ij*_ suggests that bin *i* and bin *j* are more likely to interact with each other than the other genomic loci pairs.

#### Step C. Identification of loop candidates

To minimize false positives in loop calling results, we defined a bin pair as a loop candidate only if it shows higher contact probability compared to both its global and local background. Specifically, we required the loop candidate to satisfy the following criteria:

1. Its average normalized contact probability of all single cells is greater than 0 (i.e., with respect to global background).
2. More than 10% of all single cells have normalized contact probability above 1.96 at the loop candidate (i.e., Z-score>1.96, corresponding to *p*-value<0.05, with respect to global background).
3. For each 10Kb bin pair (*i*, *j*), we defined its local neighborhood as all 10Kb bin pairs (*m*, *n*) such that 30Kb ≤ *max*{*d*(*i*, *m*), *d*(*j*, *n*)} ≤ 50Kb (**Supplementary Fig. 6**), where *d*(*i*, *m*) is the genomic distance between the center of bin *i* and the center of bin *m*. Here we did not consider the bin pairs within 20Kb of bin pair (*i*, *j*) as part of its local neighborhood because they can be part of the same loop cluster centered at bin pair (*i*, *j*). We then compared the normalized contact probability at bin pair (*i*, *j*) with the mean of the normalized contact probability of all 96 10Kb bin pairs within its local neighborhood region, and applied the paired *t*-test across all single cells to obtain a *p*-value. We further converted *p*-values into false discovery rates (FDRs) using the Benjamin-Hochberg procedure, again stratified by 1D genomic distance. The loop candidates must have FDR<10% and *t*-statistics greater than 3 in the paired *t*-test (i.e., with respect to local background).
4. Motivated by the HiCCUPS algorithm^14^, we also required the loop candidate to have at least 33% higher average normalized contact frequency than its circle, donut and lower left background and 20% higher average normalized contact frequency than its horizontal and vertical background (**Supplementary Fig. 6**) (i.e., with respect to local background).
5. Finally, we removed the loop candidates with either end having low mappability score (≤0.8), or overlapping with the ENCODE blacklist regions (http://mitra.stanford.edu/kundaje/akundaje/release/blacklists/mm10-mouse/mm10.blacklist.bed.gz for mm10 and https://www.encodeproject.org/files/ENCFF001TDO/ for hg19). The sequence mappability for each 10Kb bin is calculated based on our previous study^45^, and it can be downloaded from http://enhancer.sdsc.edu/yunjiang/resources/genomic_features/.

#### Step D. Clustering of loop candidates and identifying the summit(s) as final outputs

For each loop candidate (*i*, *j*), we defined its surrounding area as all 10Kb bin pairs (*m*, *n*) such that *max*{*d*(*i*, *m*), *d*(*j*, *n*)} ≤ 20Kb, where *d*(*i*, *m*) is the genomic distance between the center of bin *i* and the center of bin *m*. We defined a loop candidate as a singleton if there is no other loop candidate within its surrounding area, and removed all singletons from downstream analysis since the singletons are likely to be false positives.

To group the remaining non-singleton loop candidates into clusters, we adopted the Rodriguez and Laio’s algorithm^21^. Specifically, for each loop candidate (*i*, *j*), we first counted the number of loop candidates in its adjacent neighborhood regions: (*m*, *n*): *max*{*d*(*i*, *m*), *d*(*j*, *n*)} ≤ 10Kb, and defined this number as its local density *ρ*(*i*, *j*). Next, we calculated the minimum Euclidean distance between the loop candidate (*i*, *j*) and any other loop candidate with higher local density on the same chromosome, defined as *δ*(*i*, *j*):

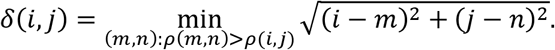

If the loop candidate (*i*, *j*) has the highest local density (i.e., *ρ*(*i*, *j*) = 9), *δ*(*i*, *j*) is defined as:

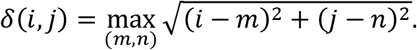

We then selected the loop candidates which have high local density *ρ*, and are relatively far away from the other loop candidates with higher local density, i.e., high *δ*, as loop cluster centers. To determine the cutoff values of *ρ* and *δ* for such centers, we implemented an algorithm similar to the ROSE algorithm^46^, which is used to identify super-enhancers. Specifically, let *ρ*_*max*_ and *δ*_*max*_ represent the maximal value of *ρ* and *δ* of all loop candidates on each chromosome, respectively. We defined *ρ*′(*i*, *j*) = *ρ*(*i*, *j*)/*ρ*_*max*_ and *δ*′(*i*, *j*) = *δ*(*i*, *j*)/*δ*_*max*_ such that both *ρ*′(*i*, *j*) and *δ*′(*i*, *j*) are within range [0,1]. We then defined *η*(*i*, *j*) = *ρ*′(*i*, *j*) * *δ*′(*i*, *j*), ordered all loop candidates by their *η* in the descending order, and plotted the rank of *η* against the value of *η*. In this plot, we selected the reflection point such that the slope at the reflection point is one. All loop candidates with *η* larger than *η* at the reflection point were chosen to be the loop cluster centers. After finding the loop cluster centers, we assigned each remaining loop candidate to the same loop cluster as its nearest neighbor with higher local density *ρ*.

Within each loop cluster, we defined the loop candidate with the lowest FDR as the first summit of the cluster. For the first summit (*i*, *j*), we defined its surrounding area as all 10Kb bin pairs (*m*, *n*) such that *max*{*d*(*i*, *m*), *d*(*j*, *n*)} ≤ 20Kb, and removed all loop candidates within its surrounding area. Next, we selected the loop candidate with the lowest FDR among the remaining ones (if there is any) as the second summit of this cluster. We then removed all loop candidates within the surrounding area of the second summit in the same way as we did for the first summit, and searched for the third summit (if there is any) with the lowest FDR among the remaining loop candidates. Such procedure was iterated until there are no loop candidates left in this cluster. Notably, one loop cluster may contain multiple summits. SnapHiC algorithm outputs a file containing the summit(s) of each loop cluster as its final chromatin loop list.

### Identification of chromatin loops with SnapHiC

We applied SnapHiC to scHi-C data from 10, 25, 50, 75, 100, 200, 300, 400, 500, 600, 700 and 742 mES cells and each of the 14 cell clusters from sn-m3C-seq data of human prefrontal cortex to call chromatin loops at 10Kb resolution between 100Kb and 1Mb region on autosomal chromosomes.

We did not take bin pairs within 100Kb into consideration because they do not have complete information in their local neighborhood (refer to “**SnapHiC algorithm**”). We also evaluated the bin pairs beyond 1Mb distance. When we extended the maximal genomic distance from 1Mb to 2Mb for loop calling using scHi-C data from 742 mES cells, only 4.6% SnapHiC-identified loops (758 out of 16,654) are between 1Mb and 2Mb. Therefore, we restricted our loop calling from 100Kb to 1Mb genomic distance for all the datasets mentioned in this study. In practice, we also suggest using 1Mb as the maximal 1D genomic distance for loop calling to save computational cost.

### Visualization of scHi-C and sn-m3C-seq data using percentage (%) of outlier cells matrix

We first computed the % of outlier cells (i.e., the proportion of cells with normalized contact probability > 1.96), and then took the integer ceiling of 100 * (% of outlier cells) to create a count matrix. We then used the Juicer^47^ software to convert the count matrix into a .hic file and visualize it in Juicebox^48^.

### Computational cost (memory, time) of SnapHiC

To assess the relationship between the number of cells and running time, we tested the running time of SnapHiC on 25, 50, 100, 200, 300 and 400 mES cells (10Kb resolution, searching for loops between 100Kb to 1Mb genomic distance) and found its running time increases linearly with the increase of cell numbers (**Supplementary Fig. 7**).

As described in our GitHub website (https://github.com/HuMingLab/SnapHiC), SnapHiC consists of two steps: (1) applying the random walk with restart (RWR) algorithm to impute contact probability within each single cell, and (2) integrating imputed contact probability matrices from all single cells to identify significant chromatin loops. Since the RWR algorithm can be applied to each chromosome in each single cell in parallel, in step 1, using as many processors as possible (e.g., maximal N = # of cells * # of chromosomes) can speed up the computation. Resolution and chromosome size are two important factors to determine the required memory per processor in step 1. For human or mouse genome at 10Kb resolution, we recommend allocating at least 30GB of memory for each processor. In the benchmarking experiments shown in **Supplementary Fig. 7**, we used 45 processors (15 nodes, 3 processors per node) for step 1, where each node has 96GB of memory, and it takes around 2.4 hours to process 100 cells.

In step 2, since the computation is performed jointly for all cells and separately for each chromosome, we recommend using the same number of processors as the number of chromosomes. Using more processors than that will be a waste of computing resources. It is also important to ensure that each processor has access to sufficient memory for the computation over all cells, and the amount of memory needed is correlated with the range of 1D genomic distance, the bin resolution, and to a less extent to the number of cells. Increasing the number of cells, slightly adds to the memory usage, however, since we only load the indices in the matrix that are used in each step of the computation, this increase in memory usage is sublinear in regard to the increase in the number of cells. In the benchmarking experiments shown in **Supplementary Fig. 7**, we used 20 processors (5 nodes, 4 processors per node) for step 2, where each node has 96GB of memory, and it takes around 0.7 hours to process 100 cells in step 2.

### Generation of aggregated contact matrix for scHi-C and sn-m3C-seq data

We pooled contacts from single cells of interest to create the aggregated contact matrix in .hic format using Juicer with KR normalization^47^. Only intra-chromosomal contacts >2Kb away are used.

### Identification of HiCCUPS loops from aggregated contact matrix

We applied the HiCCUPS^14^ to the aggregated contact matrix after pooling the contacts from single cells of interest and calling loops at 10Kb resolution with the following parameters: “--ignore_sparsity −r 10000 −k KR −f.1 −p 2 −i 5 −t 0.02,1.5,1.75,2 −d 20000”. Due to the sparsity of the aggregated contact matrix generated using single cell data, KR normalization may not always converge. Therefore, for some datasets, no HiCCUPS loops can be identified on specific chromosomes where KR-normalized matrices are not available.

To ensure a fair comparison of HiCCUPS-identified loops with SnapHiC-identified loops, we further filtered the HiCCUPS-identified loops by selecting the intra-chromosomal ones within genomic distance 100Kb~1Mb and removing the loops whose anchor bins have low mappability (≤0.8) or overlap with the ENCODE blacklist regions (refer to **Step C** in “**SnapHiC algorithm**”).

### Definition of loop overlap

Let bin pair (*i*, *j*) represent a loop in set *A*. We define it overlaps with a loop in set *B*, if and only if there exists a loop (*m*, *n*) in set *B* such that max(*d*_*im*_, *d*_*jn*_) ≤ 20Kb, where *d*_*im*_ is the 1D genomic distance between the middle base pair of bin *i* and the middle base pair of bin *m*. We allow up to 20Kb gap in the definition of loop overlap, since SnapHiC outputs summits, and bin pairs within 20Kb of the summit can be part of the same loop cluster.

### Sub-sampling of scHi-C and sn-m3C-seq data

For scHi-C data from mES cells, we randomly permuted the order of all 742 cells, and selected the first 10, 25, 50, 75, 100, 200, 300, 400, 500, 600, 700 cells from all 742 cells to create a series of sub-sampled datasets. Notably, the dataset with fewer cells is always a subset of the dataset with more cells.

For sc-m3C-seq data from human prefrontal cortex, we randomly permuted the order of all 338 astrocytes, 323 microglia and 1,038 oligodendrocytes and selected the first 261 astrocytes, microglia and oligodendrocytes to create the sub-sampled datasets for astrocytes, microglia and oligodendrocytes, respectively.

### Reproducibility of SnapHiC- and HiCCUPS-identified loops

Suppose we have two sets of loop list *A* and *B*. Let *P*_*A*_ represent the proportion of loops in set *A* overlapped with loops in set *B* (up to 20Kb gap, see **Definition of loop overlap**) and let *P*_*B*_ represent the proportion of loops in set *B* overlapped with loops in set *A*. We used (*P*_*A*_ + *P*_*B*_)/2 to measure the reproducibility of loops in the two sets.

To test the reproducibility of SnapHiC and HiCCUPS, we first randomly split all 742 mES cells into two groups where each group consists of 371 cells, and then applied SnapHiC and HiCCUPS to identify loops for each group. The reproducibility of SnapHiC- and HiCCUPS-identified loops between two sets of 371 cells are calculated as described above. We repeated such random splitting and loop calling analysis ten times, and reported the mean of reproducibility of SnapHiC- and HiCCUPS-identified loops. We further used the paired *t*-test to evaluate the statistical significance of the difference in reproducibility between these two methods.

### Generation of the reference loop lists for calculation of precision, recall and F1 score

For mES cells, the HiCCUPS loops at 10Kb resolution from bulk *in situ* Hi-C data were called as previously described^23^ using the pooled datasets of all 4 biological replicates from Bonev *et al*. study^22^. MAPS pipeline was applied to H3K4me3 PLAC-seq data^23^, cohesin HiChIP data^24^ and H3K27ac HiChIP data^25^ to call significant interactions at 10Kb resolution within 1Mb genomic distance. We combined the above four loop lists and further filtered by selecting the intra-chromosomal loops within genomic distance 100Kb~1Mb and removing loops where anchor bins have low mappability (≤0.8) or overlap with the ENCODE blacklist regions to create the final reference loop list (**Supplementary Table 4**).

For oligodendrocytes, microglia and eight neuronal subtypes from human prefrontal cortex, we used MAPS-identified interactions from H3K4me3 PLAC-seq data of purified oligodendrocytes, microglia and neurons as their reference loop list, respectively (provided in Supplementary Table 5 in Nott et al. study^29^). We first filtered the list by selecting the intra-chromosomal loops with genomic distance 100Kb~1Mb and removing loops where anchor bins have low mappability (≤0.8) or overlap with the ENCODE blacklist regions. We further selected the loops in which at least one end contains active promoters of the corresponding cell type to create the final reference loop list (**Supplementary Table 9**).

### Calculation of precision, recall and F1 score

Let *N* represent the number of loops in the reference loop list for the cell type of interest. Suppose SnapHiC (or HiCCUPS) identifies *M* loops from the same cell type, and *m* of them overlapped with loops in the reference loop list (see **Definition of loop overlap**). The precision is calculated as *m*/*M*. Suppose among all *N* loops in the reference loop list, *n* loops overlapped with SnapHiC-(or HiCCUPS-) identified loops. The recall is calculated as *n*/*N*. Notably, *m* and *n* may not be equal since we allow up to a 20Kb gap between two overlapped loops. The F1 score is the harmonic mean of the precision and recall and is calculated as below:

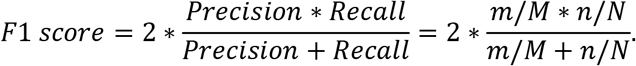

For mES cells, we used all SnapHiC- or HiCCUPS-identified loops for the above calculation. For oligodendrocytes, microglia and eight neuronal subtypes, we only selected the SnapHiC- or HiCCUPS-identified loops in which at least one anchor contains active promoters of the corresponding cell type for this calculation, since the available reference loop lists are called from H3K4me3 PLAC-seq data, which can only detect interactions centered at promoter regions.

### Aggregate peak analysis (APA)

We used the Juicer^47^ software with the command “java -jar juicer_tools_1.19.02.jar apa −r 10000 -k KR −u input.hic loops.txt APA” to perform the aggregate peak analysis. We reported “P2LL” (also known as the APA score) and “ZscoreLL” to evaluate the enrichment of SnapHiC-identified loops with respect to the lower left background.

### CTCF motif orientation analysis

We obtained the CTCF ChIP-seq peaks of mES cells from a previous study^49^, and used FIMO^50^ with default parameters and the CTCF motif (MA0139.1) from the JASPAR^51^ database to search for CTCF sequence motifs among those CTCF ChIP-seq peaks. Based on this CTCF motif list, we then selected a subset of testable SnapHiC-identified loops in which both ends contain either a single CTCF motif or multiple CTCF motifs in the same direction. Finally, we calculated the proportion of convergent, tandem and divergent CTCF motif pairs among all testable loops.

### Visualization of CTCF and H3K27ac ChIP-seq data from mES cells

We downloaded the signal tracks from the ENCODE portal^33,52^ (https://www.encodeproject.org/) with the following identifiers: ENCFF230RNU (for H3K27ac) and ENCFF069PTO (for CTCF) for **Fig. 1d**.

### Definition of cell-type-specific SnapHiC loops

We used the SnapHiC loops identified from sub-sampled astrocytes, microglia, oligodendrocytes datasets, and L2/3 excitatory neurons (all with 261 cells) to define cell-type-specific loops. Specifically, we defined a loop identified from one cell type as cell-type-specific, if it did not overlap (up to 20Kb gap, see **Definition of loop overlap**) with loops identified from any of the other three cell types.

### Selection of genes associated with cell-type-specific SnapHiC loops

We first used the Gencode v34 (GRCh37) to obtain the location of transcription start site (TSS) for 19,079 protein-coding genes in human autosomal chromosomes, and then selected genes where TSS overlaps cell-type-specific loops for astrocytes, L2/3 excitatory neurons, microglia and oligodendrocytes, respectively.

### Processing of ATAC-seq and H3K27ac ChIP-seq data from four brain cell types

The ATAC-seq and H3K27ac ChIP-seq data from human astrocytes, oligodendrocytes, microglia and neurons are from the previous study^29^ and are processed with ENCODE ATAC-seq and ChIP-seq pipelines as previously described^29^. The normalized bigwig tracks with RPKM as the *Y*-axis are generated for visualization in **Fig. 2e**.

### Processing of RNA-seq from four brain cell types

The RNA-seq data from human astrocytes, oligodendrocytes, microglia and neurons are acquired from the previous study^30^. The alignment and quantification are performed with pipeline: https://github.com/ren-lab/rnaseq-pipeline. Briefly, we first aligned RNA-seq raw reads to hg19. Next, we used Gencode GTF gencode.v19.annotation.gtf for hg19 with STAR^53^ following the ‘ENCODE’ options outlined in the STAR manual (http://labshare.cshl.edu/shares/gingeraslab/www-data/dobin/STAR/STAR.posix/doc/STARmanual.pdf). We then used Picard (http://broadinstitute.github.io/picard/) to remove PCR duplicates. We also generated the normalized bigwig tracks with RPKM (reads per kilobase of a transcript, per million mapped reads) as the *Y*-axis for visualization in **Fig. 2e**.

### Enrichment analysis of ATAC-seq or H3K27ac ChIP-seq signals at cell-type-specific loops

To quantify the intensity of ATAC-seq or H3K27ac ChIP-seq signals at cell-type-specific loops in the cell type of interest, we first calculated reads per million (CPM) values in each 10Kb anchor of the cell-type-specific loops using ATAC-seq or H3K27ac ChIP-seq data from the cell type of interest. To minimize the background noise, we only considered the reads falling into the ATAC-seq or H3K27ac ChIP-seq peak regions defined in the cell type of interest but not all the reads in the entire 10Kb bin. If there are multiple ATAC-seq or H3K27ac ChIP-seq peaks in the same 10Kb bin, we then added up the CPM values and took the sum as the value for that 10Kb bin. Since each loop has two anchors, we took their average CPM to represent the intensity of ATAC-seq or H3K27ac ChIP-seq signal for that loop in the cell type of interest. Lastly, we applied the paired Wilcoxon signed-rank test on log_2_(CPM+1) values from different combinations of cell types of interest and the cell-type-specific loop sets to test whether there is a significantly difference (**Fig. 2c**).

### Gene expression analysis at cell-type-specific loops

We obtained the FPKM values of each protein-coding genes in human astrocytes, neurons, microglia and oligodendrocytes from Supplementary Table 4 provided in the previous study (Col P-U for astrocytes, Col AB for neurons, Col AC-AG for oligodendrocytes, and Col AH-AJ for microglia in the “Human data only” tab)^30^. For each gene, we took the average of FPKM across biological replicates of the same cell type. For the selected genes where promoters are overlapped with cell-type-specific loops, we applied the Wilcoxon signed-rank test to evaluate whether they are highly expressed in the matched cell type.

### Gene ontology enrichment analysis

We used Metascape^31^ to perform gene ontology enrichment analysis for selected genes where promoters overlapped with cell-type-specific loops, and reported the top seven enriched biological processes.

## References

1 Zheng, H. & Xie, W. The role of 3D genome organization in development and cell differentiation. Nature reviews. Molecular cell biology 20, 535–550, doi:10.1038/s41580-019-0132-4 (2019).

2 Schmitt, A. D., Hu, M. & Ren, B. Genome-wide mapping and analysis of chromosome architecture. Nature reviews. Molecular cell biology 17, 743–755, doi:10.1038/nrm.2016.104 (2016).

3 Yu, M. & Ren, B. The Three-Dimensional Organization of Mammalian Genomes. Annu Rev Cell Dev Biol 33, 265–289, doi:10.1146/annurev-cellbio-100616-060531 (2017).

4 Nagano, T. et al. Single-cell Hi-C reveals cell-to-cell variability in chromosome structure. Nature 502, 59–64, doi:10.1038/nature12593 (2013).

5 Nagano, T. et al. Cell-cycle dynamics of chromosomal organization at single-cell resolution. Nature 547, 61–67, doi:10.1038/nature23001 (2017).

6 Stevens, T. J. et al. 3D structures of individual mammalian genomes studied by single-cell Hi-C. Nature 544, 59–64, doi:10.1038/nature21429 (2017).

7 Ramani, V. et al. Massively multiplex single-cell Hi-C. Nature methods 14, 263–266, doi:10.1038/nmeth.4155 (2017).

8 Flyamer, I. M. et al. Single-nucleus Hi-C reveals unique chromatin reorganization at oocyte-to-zygote transition. Nature 544, 110–114, doi:10.1038/nature21711 (2017).

9 Collombet, S. et al. Parental-to-embryo switch of chromosome organization in early embryogenesis. Nature 580, 142–146, doi:10.1038/s41586-020-2125-z (2020).

10 Tan, L., Xing, D., Chang, C. H., Li, H. & Xie, X. S. Three-dimensional genome structures of single diploid human cells. Science (New York, N.Y.) 361, 924–928, doi:10.1126/science.aat5641 (2018).

11 Tan, L., Xing, D., Daley, N. & Xie, X. S. Three-dimensional genome structures of single sensory neurons in mouse visual and olfactory systems. Nature structural & molecular biology 26, 297–307, doi:10.1038/s41594-019-0205-2 (2019).

12 Li, G. et al. Joint profiling of DNA methylation and chromatin architecture in single cells. Nature methods 16, 991–993, doi:10.1038/s41592-019-0502-z (2019).

13 Lee, D. S. et al. Simultaneous profiling of 3D genome structure and DNA methylation in single human cells. Nature methods 16, 999–1006, doi:10.1038/s41592-019-0547-z (2019).

14 Rao, Suhas S. P. et al. A 3D Map of the Human Genome at Kilobase Resolution Reveals Principles of Chromatin Looping. Cell 159, 1665–1680 (2014).

15 Ay, F., Bailey, T. L. & Noble, W. S. Statistical confidence estimation for Hi-C data reveals regulatory chromatin contacts. Genome Res 24, 999–1011, doi:10.1101/gr.160374.113 (2014).

16 Kaul, A., Bhattacharyya, S. & Ay, F. Identifying statistically significant chromatin contacts from Hi-C data with FitHiC2. Nature protocols 15, 991–1012, doi:10.1038/s41596-019-0273-0 (2020).

17 Xu, Z. et al. A hidden Markov random field-based Bayesian method for the detection of long-range chromosomal interactions in Hi-C data. Bioinformatics (Oxford, England) 32, 650–656, doi:10.1093/bioinformatics/btv650 (2016).

18 Xu, Z., Zhang, G., Wu, C., Li, Y. & Hu, M. FastHiC: a fast and accurate algorithm to detect long-range chromosomal interactions from Hi-C data. Bioinformatics (Oxford, England) 32, 2692–2695 (2016).

19 Li, X., An, Z. & Zhang, Z. Comparison of computational methods for 3D genome analysis at single-cell Hi-C level. Methods, doi:10.1016/j.ymeth.2019.08.005 (2019).

20 Zhou, J. et al. Robust single-cell Hi-C clustering by convolution- and random-walk-based imputation. Proceedings of the National Academy of Sciences of the United States of America 116, 14011–14018, doi:10.1073/pnas.1901423116 (2019).

21 Rodriguez, A. & Laio, A. Clustering by fast search and find of density peaks. Science (New York, N.Y.) 344, 1492–1496, doi:10.1126/science.1242072 (2014).

22 Bonev, B. et al. Multiscale 3D Genome Rewiring during Mouse Neural Development. Cell 171, 557–572.e524, doi:10.1016/j.cell.2017.09.043 (2017).

23 Juric, I. et al. MAPS: model-based analysis of long-range chromatin interactions from PLAC-seq and HiChIP experiments. PLoS computational biology In press., doi:10.1101/411835 (2019).

24 Mumbach, M. R. et al. HiChIP: efficient and sensitive analysis of protein-directed genome architecture. Nature methods 13, 919–922, doi:10.1038/nmeth.3999 (2016).

25 Mumbach, M. R. et al. Enhancer connectome in primary human cells identifies target genes of disease-associated DNA elements. Nature genetics 49, 1602–1612, doi:10.1038/ng.3963 (2017).

26 Fudenberg, G. et al. Formation of Chromosomal Domains by Loop Extrusion. Cell Rep 15, 2038–2049, doi:10.1016/j.celrep.2016.04.085 (2016).

27 Li, Y. et al. CRISPR reveals a distal super-enhancer required for Sox2 expression in mouse embryonic stem cells. PLoS One 9, e114485, doi:10.1371/journal.pone.0114485 (2014).

28 Schoenfelder, S. et al. The pluripotent regulatory circuitry connecting promoters to their long-range interacting elements. Genome Res 25, 582–597, doi:10.1101/gr.185272.114 (2015).

29 Nott, A. et al. Brain cell type-specific enhancer-promoter interactome maps and disease-risk association. Science (New York, N.Y.) 366, 1134–1139, doi:10.1126/science.aay0793 (2019).

30 Zhang, Y. et al. Purification and Characterization of Progenitor and Mature Human Astrocytes Reveals Transcriptional and Functional Differences with Mouse. Neuron 89, 37–53, doi:10.1016/j.neuron.2015.11.013 (2016).

31 Zhou, Y. et al. Metascape provides a biologist-oriented resource for the analysis of systems-level datasets. Nature communications 10, 1523, doi:10.1038/s41467-019-09234-6 (2019).

32 Lander, E. S. et al. Initial sequencing and analysis of the human genome. Nature 409, 860–921, doi:10.1038/35057062 (2001).

33 Consortium, E. P. An integrated encyclopedia of DNA elements in the human genome. Nature 489, 57–74, doi:10.1038/nature11247 (2012).

34 Roadmap Epigenomics, C. et al. Integrative analysis of 111 reference human epigenomes. Nature 518, 317–330, doi:10.1038/nature14248 (2015).

35 Maurano, M. T. et al. Systematic localization of common disease-associated variation in regulatory DNA. Science (New York, N.Y.) 337, 1190–1195, doi:10.1126/science.1222794 (2012).

36 Shen, Y. et al. A map of the cis-regulatory sequences in the mouse genome. Nature 488, 116–120, doi:10.1038/nature11243 (2012).

37 Andersson, R. et al. An atlas of active enhancers across human cell types and tissues. Nature 507, 455–461, doi:10.1038/nature12787 (2014).

38 Jansen, I. E. et al. Genome-wide meta-analysis identifies new loci and functional pathways influencing Alzheimer's disease risk. Nature genetics 51, 404–413, doi:10.1038/s41588-018-0311-9 (2019).

39 Demontis, D. et al. Discovery of the first genome-wide significant risk loci for attention deficit/hyperactivity disorder. Nature genetics 51, 63–75, doi:10.1038/s41588-018-0269-7 (2019).

40 Grove, J. et al. Identification of common genetic risk variants for autism spectrum disorder. Nature genetics 51, 431–444, doi:10.1038/s41588-019-0344-8 (2019).

41 Stahl, E. A. et al. Genome-wide association study identifies 30 loci associated with bipolar disorder. Nature genetics 51, 793–803, doi:10.1038/s41588-019-0397-8 (2019).

42 Savage, J. E. et al. Genome-wide association meta-analysis in 269,867 individuals identifies new genetic and functional links to intelligence. Nature genetics 50, 912–919, doi:10.1038/s41588-018-0152-6 (2018).

43 Howard, D. M. et al. Genome-wide meta-analysis of depression identifies 102 independent variants and highlights the importance of the prefrontal brain regions. Nature neuroscience 22, 343–352, doi:10.1038/s41593-018-0326-7 (2019).

44 Pardinas, A. F. et al. Common schizophrenia alleles are enriched in mutation-intolerant genes and in regions under strong background selection. Nature genetics 50, 381–389, doi:10.1038/s41588-018-0059-2 (2018).

45 Hu, M. et al. HiCNorm: removing biases in Hi-C data via Poisson regression. Bioinformatics (Oxford, England) 28, 3131–3133, doi:10.1093/bioinformatics/bts570 (2012).

46 Whyte, W. A. et al. Master transcription factors and mediator establish super-enhancers at key cell identity genes. Cell 153, 307–319, doi:10.1016/j.cell.2013.03.035 (2013).

47 Durand, N. C. et al. Juicer Provides a One-Click System for Analyzing Loop-Resolution Hi-C Experiments. Cell systems 3, 95–98, doi:10.1016/j.cels.2016.07.002 (2016).

48 Durand, N. C. et al. Juicebox Provides a Visualization System for Hi-C Contact Maps with Unlimited Zoom. Cell systems 3, 99–101, doi:10.1016/j.cels.2015.07.012 (2016).

49 Kubo, N. et al. CTCF Promotes Long-range Enhancer-promoter Interactions and Lineage-specific Gene Expression in Mammalian Cells. 2020.2003.2021.001693, doi:10.1101/2020.03.21.001693 %J bioRxiv (2020).

50 Grant, C. E., Bailey, T. L. & Noble, W. S. FIMO: scanning for occurrences of a given motif. Bioinformatics (Oxford, England) 27, 1017–1018, doi:10.1093/bioinformatics/btr064 (2011).

51 Khan, A. et al. JASPAR 2018: update of the open-access database of transcription factor binding profiles and its web framework. Nucleic acids research 46, D260–d266, doi:10.1093/nar/gkx1126 (2018).

52 Davis, C. A. et al. The Encyclopedia of DNA elements (ENCODE): data portal update. Nucleic acids research 46, D794–d801, doi:10.1093/nar/gkx1081 (2018).

53 Dobin, A. et al. STAR: ultrafast universal RNA-seq aligner. Bioinformatics (Oxford, England) 29, 15–21, doi:10.1093/bioinformatics/bts635 (2013).

